# Temperature impacts on dengue incidence are nonlinear and mediated by climatic and socioeconomic factors

**DOI:** 10.1101/2022.06.15.496305

**Authors:** Devin Kirk, Samantha Straus, Marissa L. Childs, Mallory Harris, Lisa Couper, T. Jonathan Davies, Coreen Forbes, Alyssa-Lois Gehman, Maya L. Groner, Christopher Harley, Kevin D. Lafferty, Van Savage, Eloise Skinner, Mary O’Connor, Erin A. Mordecai

**Affiliations:** Department of Biology, Stanford University, Stanford, CA 94305 USA; Department of Zoology, University of British Columbia, Vancouver, BC, V6T 1Z4 Canada; Emmett Interdisciplinary Program in Environment and Resources, Stanford University, Stanford, CA 94305 USA; Departments of Botany and Forest & Conservation Sciences, University of British Columbia, Vancouver, BC, V6T 1Z4, Canada; Hakai Institute, End of Kwakshua Channel, BC, Canada; Institute for the Oceans and Fisheries, University of British Columbia, Vancouver, BC, V6T1Z4 Canada; Bigelow Laboratory for Ocean Sciences, East Boothbay, ME, 04544, USA; U.S. Geological Survey, Western Ecological Research Center at UCSB Marine Science Institute, Santa Barbara, California, USA; Department of Ecology and Evolutionary Biology and Department of Biomathematics, University of California, Los Angeles, California 90024 USA

**Keywords:** mosquito, dengue fever, climate change, thermal biology, R0, meta-analysis, weather, disease, parasite, model

## Abstract

Temperature can influence mosquito-borne diseases like dengue. These effects are expected to vary geographically and over time in both magnitude and direction and may interact with other environmental variables, making it difficult to anticipate changes in response to climate change. Here, we investigate global variation in temperature–dengue relationship by analyzing published correlations between temperature and dengue and matching them with remotely sensed climatic and socioeconomic data. We found that the correlation between temperature and dengue was most positive at intermediate (near 24°C) temperatures, as predicted from the thermal biology of the mosquito and virus. Positive temperature–dengue associations were strongest when temperature variation and population density were high and decreased with infection burden and rainfall mean and variation, suggesting alternative limiting factors on transmission. Our results show that while climate effects on diseases are context-dependent they are also predictable from the thermal biology of transmission and its environmental and social mediators.

## INTRODUCTION

Some infectious diseases are sensitive to changes in temperature (Lafferty 2009; Rohr *et al*. 2011; Altizer *et al*. 2013; Lafferty & Mordecai 2016). This is most likely for pathogens transmitted from ectothermic hosts or vectors and/or temperature-sensitive infectious stages in the environment (Molnár *et al*. 2017). The dynamics and distributions of many diseases are predicted to alter with climate change (IPCC AR6 WG2 report 2022), with effects on human illness (Zhou *et al*. 2004), food security (Chakraborty & Newton 2011), and wildlife conservation (Cohen *et al*. 2019). Accurately predicting the effects of temperature change on infectious diseases requires understanding the impact of nonlinearity and the other factors that mediate the impact of temperature on disease systems.

Temperature can increase or decrease biological rates and processes related to disease transmission depending on the context; this type of nonlinearity makes predicting changes in infectious disease with climate change difficult. The functional traits of organisms that contribute to disease transmission—such as rates of development, activity, and fecundity and probabilities of survival and reproduction—typically have hump-shaped responses to temperature, increasing from zero at a critical thermal minimum up to an optimal temperature then declining to zero at a critical thermal maximum (i.e., thermal performance curves; Angilletta 2009; Dell *et al*. 2011; Amarasekare & Savage 2012; Mordecai *et al*. 2019). As a result, population and community-level processes, including population dynamics (Savage *et al*. 2004), disease transmission (Molnár *et al*. 2013), and trophic interactions (O’Connor *et al*. 2011; Dell *et al*. 2014), also tend to respond nonlinearly to temperature, integrating influences of temperature on multiple life stages and organisms. Thus, the observed effects of temperature on ecological processes can appear idiosyncratic, changing in direction and magnitude and becoming more or less apparent under differing circumstances, belying general predictions for how ecosystems respond to climate change (Hoos & Harley 2021).

To understand this apparent context-dependence in how temperature affects disease transmission, it may be beneficial to consider the temperature–disease relationships at a more local scale. Rather than considering a full nonlinear response of disease transmission across a large temperature range, we can instead consider the rate of change in disease with respect to temperature—which may vary across ecological settings—to link locally-determined relationships across places and times. For example, we may expect that local temperature– disease relationships will be weak at the cold end of a thermal performance curve describing disease transmission or incidence versus temperature, strongly positive where the slope of the curve is highest, and zero or weak at the optimal temperature of the curve. Whether temperature increases, decreases, or has no effect on disease transmission is therefore predicted to depend on the local average temperature and its range.

In addition to the direct effects of temperature on disease transmission, other climatic and non-climatic factors may mediate local temperature effects. For example, factors such as rainfall, drought, snow, humidity, or local variability in temperature may interact with or modify the impacts of mean temperature on disease. In particular, because body temperature and water regulation are tightly linked organismal processes, rainfall and temperature often jointly determine habitat availability and organismal performance (Rozen-Rechels *et al*. 2019), as has been shown for juvenile Ixodid ticks when seeking hosts (Berger *et al*. 2014; Leal *et al*. 2020). Local temperature variability can also play a mediating role via nonlinear averaging, in which organismal performance, and resulting population and community processes at realized temperatures, differ from what would be predicted at constant mean temperatures (Paaijmans *et al*. 2009, 2010; Bernhardt *et al*. 2018). Notably, more variable temperature tends to rescue disease transmission when it is cold and impair transmission when it is warm. Beyond climatic effects, socioeconomic and anthropogenic factors impact ecological systems through processes such as land conversion, wildlife trade and consumption, and the introduction of invasive species, which drive shifts in biodiversity, resource availability, and species distributions (Mack *et al*. 2000; Krkošek *et al*. 2007; Hendershot *et al*. 2020; Glidden *et al*. 2021). For diseases, these and other socioeconomic factors such as vector control, hygiene, and healthcare can alter the suitability of a location for disease transmission and opportunities for contacts among hosts and/or vectors. Effects of temperature on disease are most detectable when conditions are otherwise suitable for transmission, and may be dampened when other key requirements like host and vector presence and contact are not met.

The net effects of nonlinearity and other factors mediating temperature impacts on disease will have consequences for human health, especially for vector-borne diseases. In particular, dengue is a climate-sensitive, tropical and subtropical disease caused by a flavivirus (DENV) primarily transmitted by female *Aedes aegypti* mosquitoes; it causes 100-400 million cases every year (WHO 2021) and cases have been increasing dramatically both regionally and globally over the last three decades (Stanaway *et al*. 2016). Notably, since mosquitoes and the parasites they harbor are ectotherms, temperature can influence multiple stages of the mosquito life cycle and transmission cycle, affecting the distribution and dynamics of disease (Liu-Helmersson *et al*. 2014; Morin *et al*. 2015; Wesolowski *et al*. 2015; Mordecai *et al*. 2017). Previous research has used a combination of experiments and mathematical modeling to first isolate the effects of temperature on different mosquito and pathogen traits (e.g., DENV development rate within the mosquito, mosquito lifespan and fecundity) and then combine these processes to understand how potential transmission rates vary across temperature (Lambrechts *et al*. 2011; Liu-Helmersson *et al*. 2014; Wesolowski *et al*. 2015; Huber *et al*. 2018; Caldwell *et al*. 2021). This has provided specific predictions for how temperature affects dengue transmission in the field: small increases in temperature should increase transmission up to the optimal temperature of 29°C, after which increases in temperature should decrease transmission (Mordecai *et al*. 2017). The greatest relative increase in transmission per degree increase in temperature is expected to occur near 25°C (i.e., the temperature at which the slope of the transmission versus temperature curve is steepest). Although some empirical support for these predictions exists at broad spatial scales in the field (e.g., Wesolowski *et al*. 2015; Mordecai *et al*. 2017; Peña-García *et al*. 2017; Caldwell *et al*. 2021), recognition of the importance of nonlinear effects of temperature on transmission, especially at local scales, remains limited.

Here, we consider dengue as a case study to examine correlations between temperature and disease transmission. Previous work has reported both positive and negative relationships between temperature and dengue outbreaks (Caldwell *et al*. 2021). We hypothesized that nonlinear effects of temperature, mediated by other climatic and non-climatic factors, might explain apparent differences in the inferred effects of temperature on dengue transmission. We searched the literature to test whether dengue transmission—measured as empirical correlations—changes nonlinearly with average study temperature and peaks near 25°C, the temperature where the slope of the transmission versus temperature curve was suggested to be greatest in a previously published trait-based mathematical model (Mordecai *et al*. 2017). We also test our predictions that the strength of correlations increase positively with temperature variation since it should be easier to detect effects of temperature when it is more variable, and either increase or decrease with precipitation mean and variability depending on whether local vector abundance is rain-or drought-driven (Lowe *et al*. 2021). Finally, we test whether correlations decrease or become more negative with infection burden in the area due to depletion of susceptible hosts, increase with population density due to larger epidemic potential, and either decrease or become more negative with income (measured as per-capita gross domestic product; GDP), which reduces outbreak potential and dampens the effects of suitable temperatures.

## METHODS

### Overview

To test our predictions, we synthesized published evidence of temperature–dengue relationships using a systematic literature review. We compiled reported correlations between temperature and dengue from previously published studies. We did not consistently have access to the underlying temperature and dengue data used in the original studies that would have allowed for a reanalysis of the raw data across locations. Instead, we paired each reported correlation with climate reanalysis data and data on factors such as wealth and human density. This means that while we did not have the underlying data used to estimate correlations in each study, we did have estimates of the average temperature, average variability in temperature and precipitation, population density, and other socioeconomic and climatic factors in each focal study area and time period.

We used this database to answer two questions: 1) Does average study temperature impact temperature–dengue relationships? and 2) How do other climatic and socioeconomic factors explain variation in temperature–dengue relationships? Below, we detail the database construction as well as the two separate analyses used to answer these questions.

### Database construction

We downloaded abstracts and study metadata (N = 454) from Web of Science on January 28, 2021 (accessed through the University of British Columbia library), using the search term TS = ((“Aedes” OR “dengue”) AND (“temperature” OR “climat*”) AND (“disease*”) AND (“model*”) AND (“incidence” or “prevalence” or “case*” or “notification*”)). We then systematically conducted several rounds of scoring to exclude studies with irrelevant or missing information. First, we read each abstract and scored it as included or excluded based on the mention of factors such as measured climatic variables and measured disease burden, incidence, or prevalence. Studies were excluded if the abstract mentioned forecasting or simulations only. In total, 189 of 454 abstracts were accepted. Next, we read each study with an accepted abstract, and scored the study as either included or excluded based on the presence of effect sizes or correlations comparing the effects of measured temperature metrics and measured disease metrics. We excluded a study if only forecasting or simulation models were presented. For this step, 95 of 189 papers with accepted abstracts were accepted.

We initially planned to collect data from studies that reported either a correlation between temperature and dengue or a coefficient estimating the effect of temperature on dengue from a regression analysis. However, our systematic literature review revealed that most of the studies using regressions incorporated different covariates into their models, ranging from accounting for no covariates to accounting for the effects of multiple temperature metrics, precipitation, GDP, and others. We conducted simulations that illustrated how these different underlying models can lead to significantly different estimates of the effects of temperature on dengue despite the temperature and dengue data remaining identical across models (Supporting Information), making comparisons across regression models unreliable for the purposes of our study. Instead, we focused all subsequent analyses on reported correlations between temperature and dengue as these models do not include any covariates, resulting in 358 reported correlations from 38 studies (Table S1).

We included methodological information (hereafter referred to as study factors) for each correlation, such as the location of the study, dates and length of the study, the types of temperature (e.g., minimum weekly temperature, mean daily temperature) and disease metrics (e.g., cases, incidence) used in the analysis, the type of correlation (Pearson, Spearman, or cross-correlation), and the temporal lag of the effect of temperature. We also complemented our database with data (hereafter referred to as extracted predictors) obtained from several other sources. We used Google Earth Engine (Gorelick *et al*. 2017) to extract information on population density and climate over the period of each study. Population density was obtained from the Global Human Settlement Population Grid (JRC 2015), using the year closest to the median year of each study period. Average daily mean air temperature, standard deviation in daily mean air temperature, mean daily precipitation, and standard deviation in daily precipitation were obtained from ERA5 (C3S 2017) and calculated over each full study period. Study locations on the scale of a single city or smaller were specified using a 5 kilometer buffer around point coordinates, while larger areas were mapped using shapefiles obtained from the Database of Global Administrative Areas (GADM 2021). To reflect the climatic and population factors most relevant to where people live (and thus where dengue cases occur), we weighted these measures over space by population density. The estimated infection burden of dengue at the country level (in the year 2010) was extracted from Bhatt *et al*. (2013) as a proxy for the degree of population immunity or susceptibility. Country level population size in 2010 and GDP per capita (adjusted for purchasing price parity in the year 2015) were obtained from the World Bank (2022). Estimated dengue incidence in 2010 was calculated as estimated burden / population size (see Supporting Information for more detail).

### Does average study temperature impact temperature–dengue effects?

To test for a relationship across studies between mean study temperature (calculated as mean average daily temperature across the study period) and observed correlation between temperature and dengue within that study, we fit a series of linear mixed effects models using reported correlations as response variables. Prior to fitting these models, we limited the dataset to exclude observations that were generated using lags > 4 months (our estimate of the maximum biologically relevant window on which temperature could directly affect dengue transmission), and included only one observation per location and temperature metric per study to avoid having multiple observations estimated across different lags. For example, if a study reported five correlation values between minimum monthly temperature and dengue for a specific location using lags of 0, 1, 2, 3, and 4 months, we would only select the observation closest to the midpoint of 2 months. This resulted in 78 correlation observations from 37 studies (one of the 38 studies used only lags > 4 months).

We aimed to test whether the measured relationship between temperature and dengue depended on the average temperature during the study, as well as whether ecological theory based on a lab-parameterized, trait-based model of dengue transmission across temperature (Mordecai *et al*. 2017) could accurately predict how correlations vary across mean temperature. Specifically, we fit a null model and four alternative mixed-effects models in R (R Core Team 2021) using maximum likelihood with the *lmer* function (Bates *et al*. 2015). The null model included only a random effect for study ID, the basic model included the study ID random effect and an additional fixed effect for the type of temperature metric used in the study (minimum, mean, or maximum temperature), and the final three models included the previously described effects and additionally a fixed effect for either a linear effect of average temperature, a quadratic effect of average temperature, or for the derivative of the transmission curve from Mordecai *et al*. (2017) evaluated at the mean study temperature. The purpose of including this final model was to compare the observed relationship based on reported correlations to the a priori theoretical relationship that first motivated us to look for a concave-down pattern in correlations between 20°C and 29°C. However, we note that the derivative of model-predicted dengue basic reproduction number (R_0_) represents a mathematical quantity that is distinct from a correlation between temperature and dengue. Therefore, while we suspected that these two values may follow the same qualitative patterns across temperature, they are not mathematically equivalent because R_0_ does not predict incidence directly (Smith *et al*. 2007).

We compared the five models using AIC (R Core Team 2021) and extracted Nagelkerke’s pseudo-R^2^ values using the MuMIn package (Bartoń 2020). We did not incorporate error around reported correlation estimates because this information was not available, though we repeated the analyses described here while weighting estimates by the square root of their sample size, a method used in meta-analyses when error estimates are unavailable (Hargreaves *et al*. 2020).

### How do other climatic and socioeconomic factors explain variation in temperature–dengue effects?

Next, we aimed to test how additional climatic factors such as precipitation and socioeconomic factors such as country-level GDP impacted the observed effects of temperature on dengue. As described in the Introduction, we predicted that temperature–dengue correlations would be more positive with higher temperature variation and population density, lower with higher infection burden and GDP, and modified (either positively or negatively) by precipitation mean and variability. While we originally intended to estimate how each of these extracted predictors separately mediates the effects of temperature, this was not possible due to the high collinearity between predictors (Fig. S2). We therefore conducted a two-step analysis, collapsing the variance from all predictors with a principal component analysis (PCA) and evaluating the PCA components along with study factors in linear regression models.

The PCA incorporated seven extracted predictors: log-transformed country-level GDP, country-level infection incidence, and five metrics calculated by study: log-transformed population density, mean precipitation, standard deviation of precipitation, standard deviation of temperature, and marginal temperature suitability (the derivative of the Mordecai *et al*. [2017] dengue transmission curve evaluated at the mean study temperature, as described above). We sampled the unique sets of these extracted predictors, then used the *principal* function from the *psych* package (Revelle 2021) to load the seven predictors across four principal components that were rotated using Varimax rotation. While traditional PCA typically rotates axes to explain the maximal amount of variation using the first component, Varimax rotation maximizes the sum of the variances of the squared loadings, allowing for better interpretability of which predictors are more strongly associated with which components.

We fit regressions using the full dataset of correlations (n=358) as response variables. Predictors included the four rotated components from the PCA analysis, as well as study factors to help control for variation introduced by different study methods: the temperature metric used in the study (minimum, mean, or maximum), the disease metric used in the study (incidence or cases), the temporal scale of the study (daily, weekly, monthly, or annual), and the type of correlation used in the study (Pearson, Spearman’s, or cross-correlation). We also included a term for a spline (dimension of the basis = 3) for the effect of temporal lag in months on the effect of temperature on dengue. We did not include interactions between these predictors. We fit the regressions using the *gam* function in the *mgcv* package (Wood 2011) due to the inclusion of the spline term for lags. We did not want studies that provided relatively more observations (either because they estimated effects across multiple lags, multiple temperature metrics, or multiple locations) to be overrepresented in our regression. We therefore bootstrapped 10,000 times, each time first sampling studies (n=38) with replacement and then sampling one observation within that study, until we generated a dataset equal in size to the original (n=358) to reduce overrepresentation of studies with many data points. We extracted the mean and 0.025 and 0.975 quantiles for each predictor coefficient estimate across the 10,000 bootstraps.

## RESULTS

We obtained 358 reported correlations between temperature and dengue from 38 studies, ranging from 1981 to 2017 and spanning seven global health regions (Southeast Asia, East Asia, South Asia, Central Latin America, Tropical Latin America, Oceania and Caribbean; Moran *et al*. 2012)(Fig. 1a-b). The estimates were variable with 19% negative and 81% positive (Fig. 1c).

**Figure 1.**
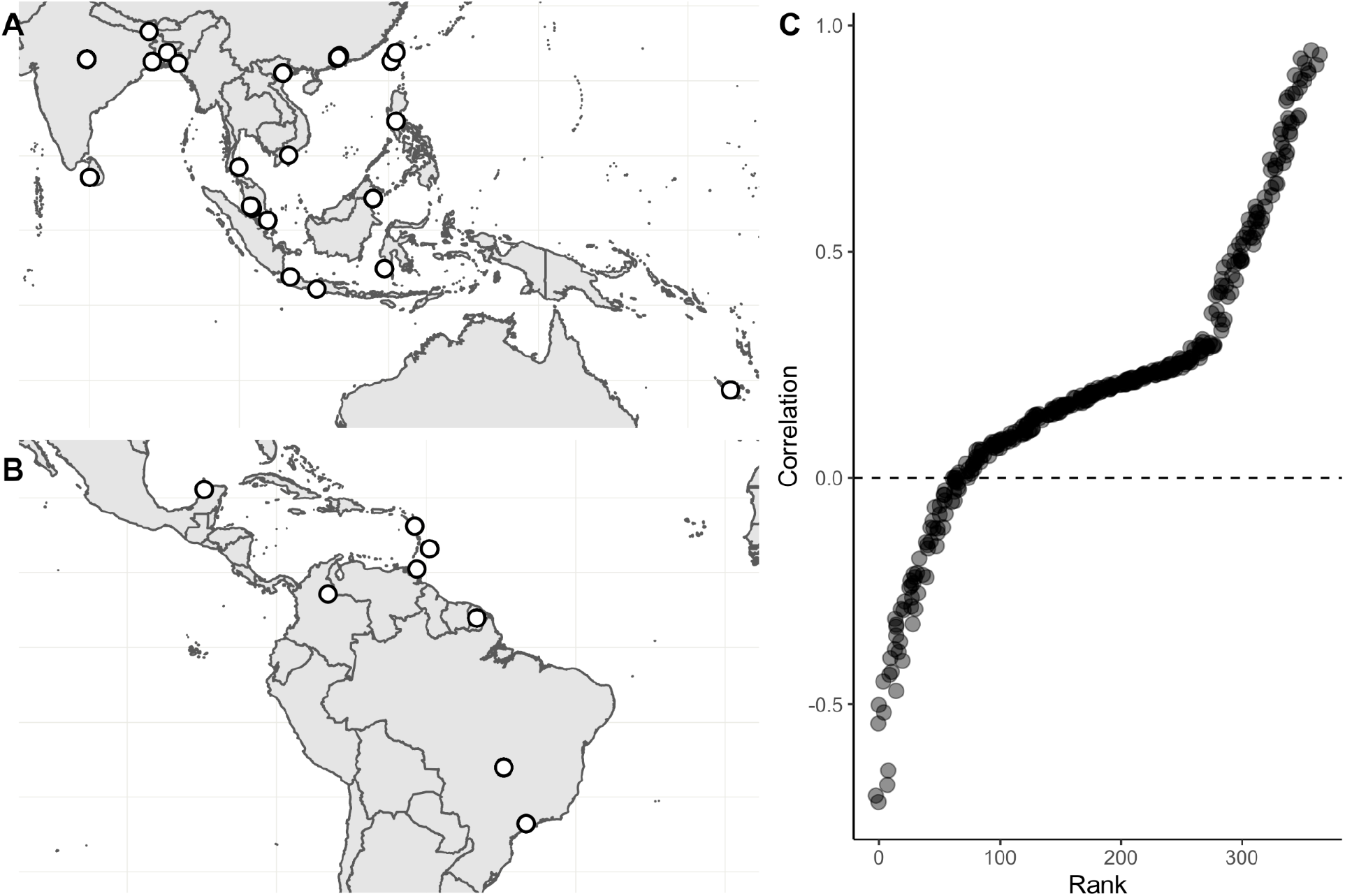
Reported correlations between temperature and dengue range from negative to positive. a) Locations of observations in the global health regions of Southeast Asia, East Asia, South Asia, and Oceania; b) Locations of observations in the global health regions of Central Latin America, Tropical Latin America, and Caribbean; c) Jittered rank-order plot of 358 reported correlations between temperature and dengue.

Supporting predictions, we found that the best model included a nonlinear (quadratic) effect of mean study temperature on reported correlations (ΔAIC from null model = 10.3; pseudo-R^2^ = 0.209). The quadratic model estimates that reported correlations peak at mean study temperatures of 24.2°C (95% CI: 23.5–24.9°C; Fig. 2). The second-best model included the nonlinear effect of mean study temperature calculated from the derivative of the Mordecai et al. (2017) transmission curve, which peaks at 25.3°C (ΔAIC = 8.0; pseudo-R^2^ = 0.165), suggesting that ecological models based on vector and parasite biology can help predict how correlations vary across average temperatures. The model incorporating a linear effect of mean study temperature (ΔAIC = 5.0; pseudo-R^2^ = 0.132) did not perform better than the basic model that did not include any effect of mean study temperature (ΔAIC = 5.9; pseudo-R^2^ = 0.119). Repeating these analyses while weighting by the square root of the study sample size produced qualitatively similar results (Supporting Information).

**Figure 2.**
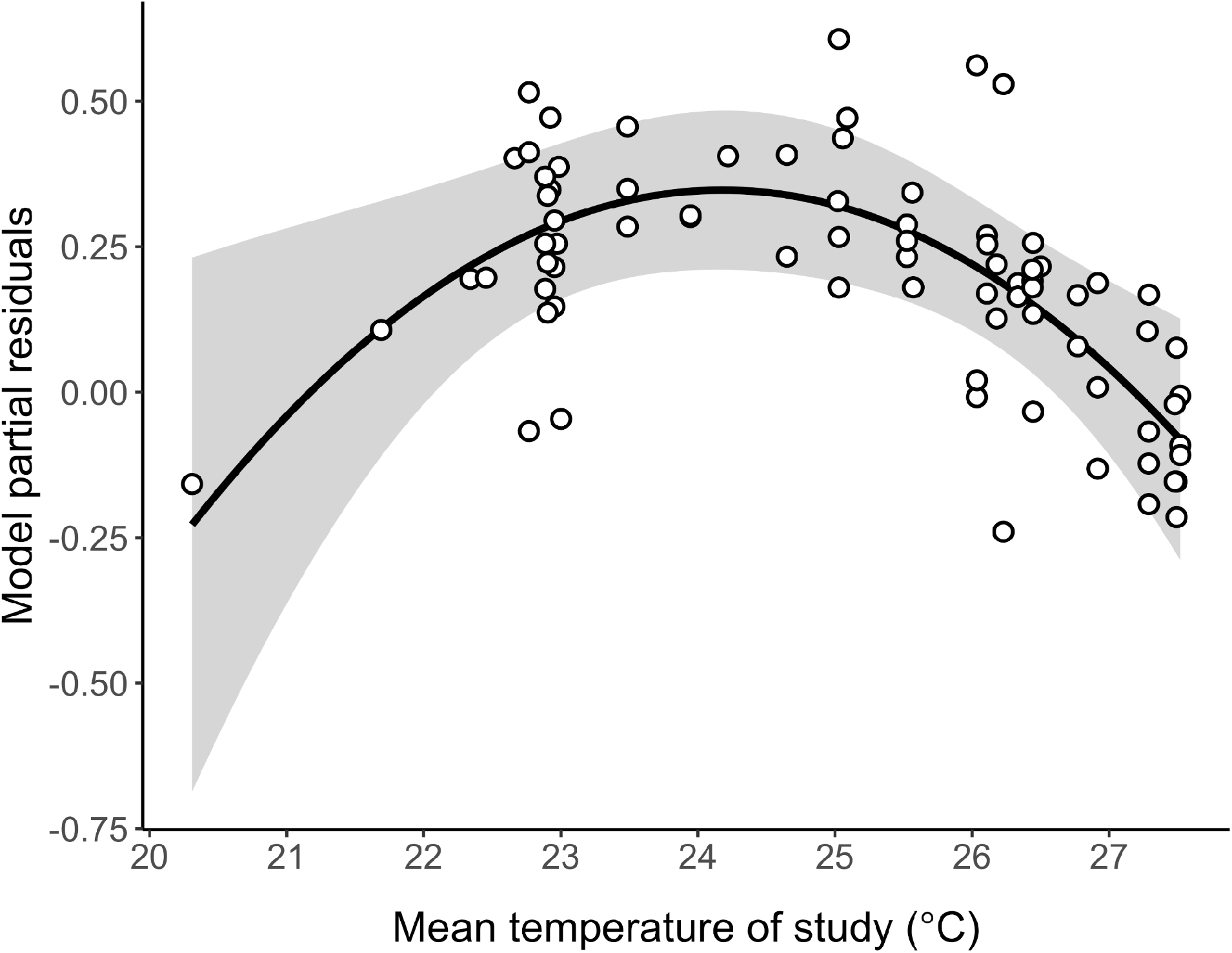
Nonlinear effects of temperature on the correlation between temperature and dengue, controlling for study factors. Quadratic model partial residuals (points) and fitted predictions (black line) with 95% confidence intervals (shaded region) for the relationship between mean study temperature and reported correlations between temperature and dengue. Partial residuals and fitted predictions are from the mixed effects model with a quadratic effect of mean study temperature (black line), which was significantly better than alternative models that included a linear effect or no effect of mean study temperature (ΔAIC from null model = 10.3; pseudo-R^2^ = 0.209). Partial residuals are calculated as model errors plus the model-estimated relationship between temperature and dengue. Confidence intervals generated using the *effects* package in R (Fox and Weisberg 2019). Figure S1 shows the same fitted model plotted over raw correlation data.

We then examined the factors beyond mean temperature that mediated the observed relationship between temperature and dengue. Using PCA to decompose correlated climatic and socioeconomic predictors into fewer, uncorrelated rotated components (RCs) meant that we were not able to estimate the specific effect of each predictor on reported relationships between temperature and dengue. However, this method was useful for identifying RCs that have significant effects on our response, which we can then interpret as the underlying predictors associated with each component having a positive or negative effect on the response.

Several RCs had a significant effect on reported correlations (Fig. 3), generally supporting our hypotheses. Infection burden (RC1) had negative effects on reported correlations, while temperature variation (RC1) and marginal temperature suitability (i.e., the derivative of the predicted transmission curve; RC3) had positive effects. We did not have a directional prediction for the effects of mean precipitation (RC2) and precipitation variation (RC2), but found that they had negative effects. Higher population density was associated with two different rotated components, and exhibited a significant, positive effect associated with RC1 and a non-significant, positive effect when combined with lower GDP (RC4). Several study factors also had significant effects on reported correlations: most notably, we found that studies that used a metric of minimum or mean temperature reported more positive correlations between temperature and dengue than those studies that used a metric of maximum temperature (Figs. S3-S4).

**Figure 3.**
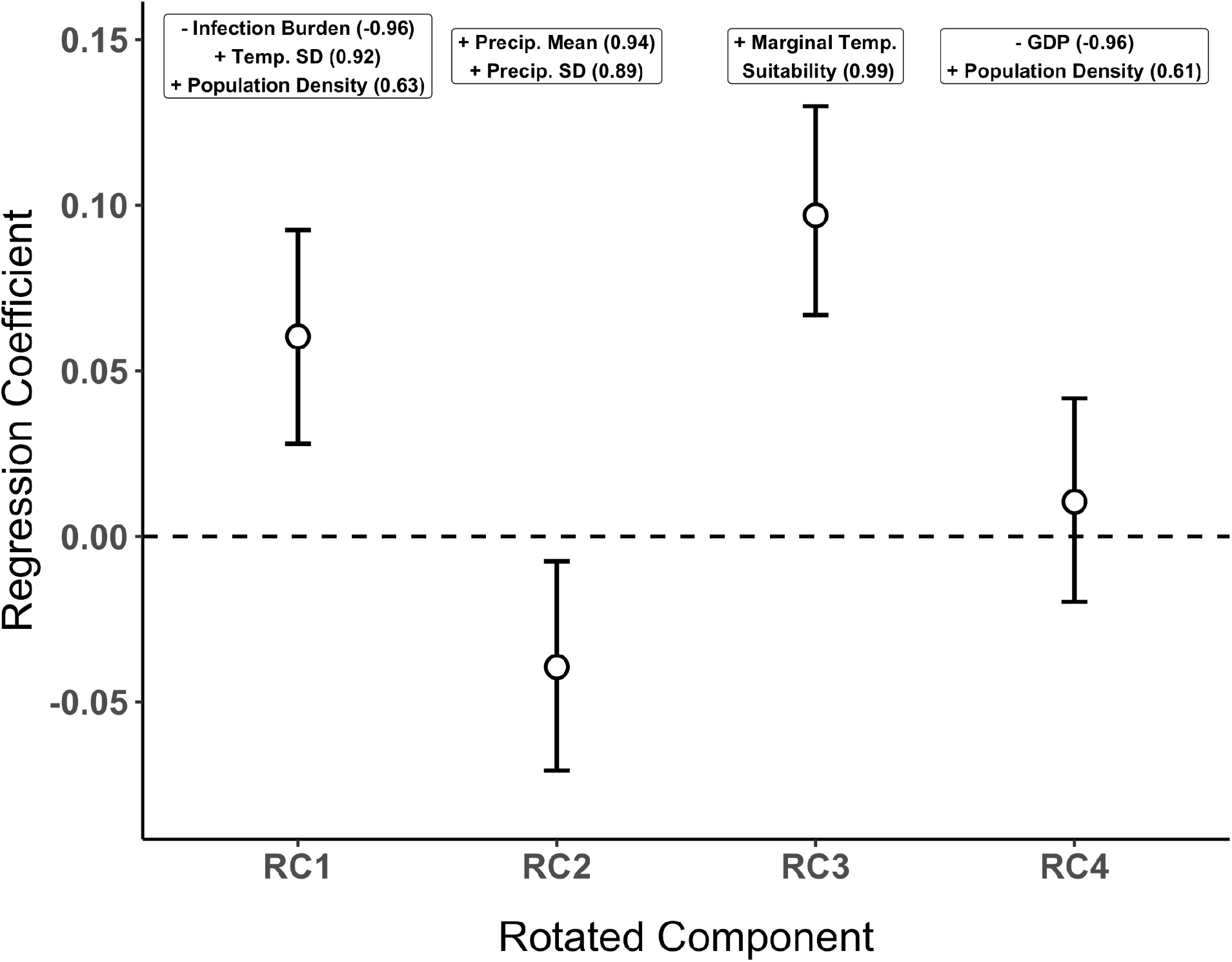
Infection burden, temperature variability, population density, precipitation, and predicted temperature suitability affect the strength of temperature – dengue correlations. Mean and 95% confidence intervals of regression coefficients for four rotated components (RC) across 10,000 bootstrap runs. Annotated text above each component lists the climatic and/or socioeconomic factors most strongly associated with that component (standardized loading > |0.6|), with +/-symbols representing the sign of the association and the numbers in parentheses representing the loading (where 1 and -1 represent the strongest positive and negative associations, respectively). The sign of each association (in boxes) combined with the signs of each respective regression coefficient (points) yields the direction of the effect of each predictor on correlations (e.g., infection burden (RC1), mean precipitation (RC2) and precipitation variation (RC2) all have significant, negative effects).

## DISCUSSION

Our examination of reported correlations between temperature and dengue support predictions that the effects of temperature on many ecological processes are nonlinear with small or negative effects expected at low and high temperatures, and large positive effects expected in some intermediate temperature range. Specifically, studies that occurred at relatively cool or warm average temperatures reported lower correlations than those that occurred at temperatures near the intermediate range, where transmission is expected to be most sensitive to temperature (24°C). Our results illustrate that locations differ in their underlying vulnerability to warming-induced disease outbreaks, and that this variability in vulnerability can be explained by nonlinearity and average temperatures, as well as other climatic and socioeconomic factors such as precipitation and disease burden.

The average temperature at which a study occurred had a significant quadratic relationship with the correlation between temperature and dengue, with a peak at 24.2°C (95% CI: 23.5–24.9°C; Fig. 2). This is close to but slightly cooler than the prediction from the derivative of the trait-based, dengue transmission curve that is informed by laboratory studies on the vector *Aedes aegypti* (25.3°C; Mordecai *et al*. 2017). Both the flexible quadratic temperature model and the *a priori* marginal temperature suitability model (the derivative of the Mordecai et al. [2017] model) were significantly better than the simpler models that assumed the correlation between temperature and dengue was constant or linear across average temperature. Further, in a PCA that controlled for multiple climatic and non-climatic factors, the component that was highly associated with the *a priori* marginal temperature suitability model had significant positive effects on correlations (Fig. 3). These results build on observations that reported temperature effects on dengue varied across average temperatures or climates (Fan *et al*. 2015; Li *et al*. 2020; Caldwell *et al*. 2021) by quantitatively testing whether effects vary nonlinearly as predicted by ecological theory. Additionally, our database and analyses differed both by using reported correlations rather than coefficients from regression models, as well as by using standardized remotely sensed temperature data across studies rather than using average temperatures reported by each original study. Overall, our results suggest that ecological theory can be used to predict how relationships between temperature and disease vary with average temperature, an often underappreciated facet of the impact of climate change on infectious disease.

In addition to temperature having a direct nonlinear impact on dengue across average study temperatures, we found that several other climatic factors mediated these effects. Mean precipitation and variation in precipitation each had significant negative effects (via their association with RC2) on reported correlations between temperature and dengue (Fig. 3). Precipitation could modulate the temperature–dengue relationship through several alternative mechanisms, though our approach does not allow us to differentiate between them. When temperature is not strongly limiting to transmission but immature vector habitat is inconsistently available, precipitation may be the main limiting factor, obscuring the relationship between temperature and dengue. Alternatively, both temperature and precipitation may be limiting in some settings, such that even when suitable temperatures occur there is insufficient vector habitat to promote transmission. Finally, correlations between temperature and rainfall regimes (e.g., seasonality) may obscure the causal relationships between each variable and dengue. While precipitation may not mediate temperature effects in all ecological or disease systems, it could play a key mediating role in systems with animals that require pools of water for habitat or breeding (e.g., other mosquito-borne diseases; Paull *et al*. 2017), in waterborne-disease systems such as cholera, and in plant systems in which rainfall has been shown to impact disease levels (McElrone *et al*. 2010; Eastburn *et al*. 2011).

In contrast to precipitation, average temperature variability during a study had significant positive effects (via its association with RC1) on the correlation between temperature and dengue, potentially because it is easier to detect correlations when temperature fluctuates over a wider range. Additionally, nonlinear averaging can cause more positive effects of temperature variation on dengue at ranges where the temperature–transmission relationship is concave-up than concave-down (Lambrechts *et al*. 2011). Consideration of temperature variability should become more important with climate change, as large changes in temperature variability and in the frequency, magnitude, and duration of temperature extremes are expected in many regions but their impacts on ecological processes have received relatively little attention (Easterling *et al*. 2000; Smith 2011; Thompson *et al*. 2013; Turner *et al*. 2020; Ma *et al*. 2021). Together these results provide an important biological insight: effects of temperature on ecological processes can be exacerbated or masked by other aspects of climate suitability, including rainfall and variation in temperature.

Immunological and other non-climatic factors also affected local relationships between temperature and dengue. As predicted, we observed a strong negative effect of infection burden (as estimated for the year 2010; Bhatt *et al*. 2013), in which locations with higher levels of dengue reported weaker or more negative correlations between temperature and dengue. One possible explanation for this is that populations with historically high dengue burden have proportionally high levels of immunity and partial immunity (Gubler 1998), thereby leaving fewer people susceptible to infection when temperature conditions become more optimal. One potential caveat when interpreting these effects is that the Bhatt *et al*. model (2013) used additional data inputs beyond dengue cases—including temperature suitability—to estimate country-level infection burden, meaning that estimated dengue burden is not completely independent from temperature. Our predictions that population density would increase temperature effects due to larger epidemic potential, while higher GDP would decrease temperature effects due to higher income leading to better health infrastructure and disease mitigation were generally supported (Fig. 3).

Because of the thermal physiology of organisms, we expect many ecological systems and processes to be nonlinearly dependent on temperature, and these temperature effects are likely to be mediated by other ecological and socioeconomic factors. Dengue provides a relatively well-studied example for detecting these nonlinear and mediated effects, which may not be possible for more data-limited ecological systems. Primary studies that investigate nonlinear effects of temperature on ecological processes explicitly, and the mediators of these effects, are critical for more generally anticipating the impact of climate change on ecological systems.

Many ecological systems are dominated by physiological processes that respond nonlinearly to temperature (Brown *et al*. 2004; Dell *et al*. 2011), making them prone to climate change impacts that vary in magnitude and direction across ecological settings. Recognizing this nonlinearity as a fundamental driver of context-dependent responses is a critical conceptual gap in many ecological studies of climate change. This can help to resolve inconsistent correlations with temperature found between different field locations, as has been found with withering syndrome in abalone (Ben-Horin *et al*. 2013) and sea star wasting disease (Eisenlord *et al*. 2016; Menge *et al*. 2016; Harvell *et al*. 2019), as well as in other ecological contexts beyond disease (Leonard 2000). At the same time, the magnitude of nonlinear effects of temperature depends on a range of environmental, anthropogenic, and biogeographic factors, including climatic variation in rainfall, temperature, humidity, and extreme events, human-driven changes in habitat structure and species composition, and evolutionary history. Together, these factors mediate ecological effects of temperature by affecting body condition, behavior, species interactions, and evolutionary processes (Huey & Kingsolver 2019). Research that combines a mechanistic understanding of the nonlinear impacts of temperature on ecological processes with explicit consideration of important modifiers of temperature responses—through either comparative approaches like that taken here or experimental approaches that manipulate multiple drivers directly (e.g., Zhu *et al*. 2016)—can help to capture realistic variation in the effects of climate change across settings.

## ACKNOWLEDGMENTS

We thank the University of British Columbia for its support for this project through the Grants for Catalyzing Research award. MJH was supported by the Knight-Hennessy Scholars Program. MLC was supported by the Illich-Sadowsky Fellowship through the Stanford Interdisciplinary Graduate Fellowship program. DK, ES, and EAM were supported by the National Institutes of Health (R35GM133439). EAM was additionally supported by the National Science Foundation (DEB-2011147, with the Fogarty International Center) and the Stanford Woods Institute for the Environment, Center for Innovation in Global Health, and King Center on Global Development.

