## Supporting Information for "Temperature impacts on dengue incidence are nonlinear and mediated by climatic and socioeconomic factors"

### Further details: Database construction

#### *Extracted predictors*

As described in the main text, we conducted a systematic literature review of studies that reported temperature effects on dengue, then matched each study with remotely sensed climate data and data on factors such as wealth and human density as potential mediators of the effects of temperature on dengue incidence.

Our database contains methodological information (i.e., study factors) for each correlation that was extracted from the studies, but we also complemented our database with data (i.e., extracted predictors) obtained from several other sources. We used Google Earth Engine (Gorelick *et al.* 2017) to extract information on population density and climate over the period of each study. Population density was obtained from the Global Human Settlement Population Grid (JRC 2015), using the year closest to the median year of each study period. Average daily mean air temperature, standard deviation in daily mean air temperature, mean daily precipitation, and standard deviation in daily precipitation were obtained from ERA5 (C3S 2017) and calculated over each full study period. Study locations on the scale of a single city or smaller were specified using point coordinates with a 5km buffer around them, while larger areas were mapped using shapefiles obtained from the Database of Global Administrative Areas (GADM)(GADM 2021).

To reflect the climatic and population factors most relevant to where people live (and thus where dengue cases occur), we weighted these measures over space by population density.

We extracted population and climatic data from 39 studies, then visually inspected the extracted mean temperature values for each location to ensure they were sensible and generally aligned with reported average climates using www.weatherspark.com. In one case, we decided to exclude a study from our database due to unreliable temperature data: our extracted mean temperature using the ERA5 dataset for the city of Cali, Colombia from 2001–2011 for one study (Eastin *et al.* 2014) was significantly lower than the average temperature reported for this city from other sources (20°C compared to >23°C reported from weatherspark, 23–24°C from WorldClim (Hijmans et al. 2005) extracted via Google Earth Engine, and 23–24°C reported in the original study). This left the final database that we used in our analyses with 38 studies comprising 358 total observations.

The estimated infection burden of dengue at the country level (in the year 2010) was extracted from Bhatt *et al.* (2013) as a proxy for the degree of population immunity or susceptibility. Data on dengue burden exists at higher spatial resolution than the country level and/or across time (compared to only in the year 2010). However, we used the data from Bhatt et al. (2013) because it represented one dataset that comprehensively covered all locations in our study, and we considered this superior to trying to compare multiple dengue datasets for different locations that were compiled using different methodologies. We note that the Bhatt *et al.* (2013) model used additional data inputs beyond dengue cases—including temperature suitability—to estimate country-level infection burden, meaning that estimated dengue burden is not completely independent from temperature. We obtained country level population size in 2010 and GDP per capita (adjusted for purchasing price parity in the year 2015) from the World Bank (2022), then calculated dengue incidence in 2010 as the estimated burden divided by population size. Most of the study locations (Table S1) could simply be matched with the name of the countries used by the World Bank to calculate GDP per capita, but we used the GDP reported for France for Guadeloupe, New Caledonia, and French Guiana and the GDP reported for China for Taiwan. Each of these four locations had population size data from 2010 specific to them.

*List of original studies used in our database*

**Table S1.** List of 38 studies in our database that report correlations between temperature and dengue, the location and years in which each study took place, and the number of observations (i.e., correlations) included in our database from each study.

| Reference | Study location | Years study took place | Number of observations |
|---|---|---|---|
| Wijaya *et al.* 2021 | Indonesia | 2009–2014 | 5 |
| Gharbi *et al.* 2011 | Guadeloupe | 2000–2007 | 2 |
| Laureano-Rosario *et al.* 2017 | Mexico | 2007–2010 | 3 |
| Minh An & Rocklöv 2014 | Vietnam | 2002–2010 | 7 |
| Wang *et al.* 2014 | China | 2000–2012 | 4 |
| Wu *et al.* 2007 | Taiwan | 1988–2003 | 12 |
| Chen *et al.* 2020 | China | 2006–2017 | 2 |
| Aswi *et al.* 2020 | Indonesia | 2013–2017 | 3 |
| Bal & Sodoudi 2020 | India | 2005–2016 | 8 |
| Oliveira *et al.* 2020 | Brazil | 2008–2015 | 22 |
| Zhu *et al.* 2019 | China | 2008–2016 | 1 |
| Jayaraj *et al.* 2019 | Malaysia | 2006–2017 | 21 |
| Ogashawara *et al.* 2019 | Brazil | 2011-–2017 | 2 |
| Chen *et al.* 2019 | China | 2014 | 26 |
| Kakarla *et al.* 2019 | India | 2010–2017 | 72 |
| Kong *et al.* 2019 | China | 2006–2014 | 12 |
| Acharya *et al.* 2018 | Nepal | 2011–2016 | 1 |
| Zahirul Islam *et al.* 2018 | Bangladesh | 2000–2009 | 2 |
| Jing *et al.* 2018 | China | 2001–2014 | 15 |

Table S1 continued.

| Reference | Study location | Years study took place | Number of observations |
|---|---|---|---|
| Withanage *et al.* 2018 | Sri Lanka | 2012–2015 | 8 |
| Carvajal *et al.* 2018 | Philippines | 2009–2013 | 3 |
| Ahmad *et al.* 2018 | Malaysia | 2014–2015 | 4 |
| Chang *et al.* 2018 | Taiwan | 2007–2011 | 3 |
| Li *et al.* 2017a | China | 1998-2014 | 12 |
| Martínez-Bello *et al.* 2017 | Colombia | 2008–2015 | 1 |
| Li *et al.* 2017b | China | 2011–2014 | 16 |
| Ramadona *et al.* 2016 | Indonesia | 2001–2013 | 1 |
| Phuong *et al.* 2016 | Vietnam | 2004–2014 | 1 |
| Teurlai *et al.* 2015 | New Caledonia | 1995–2012 | 3 |
| Liao *et al.* 2015 | Taiwan | 2004–2013 | 5 |
| Flamand *et al.* 2014 | French Guiana | 2006–2011 | 10 |
| Sang *et al.* 2014 | China | 2006–2012 | 24 |
| Wongkoon *et al.* 2013 | Thailand | 1981–2012 | 2 |
| Cheong *et al.* 2013 | Malaysia | 2008–2010 | 3 |
| Pinto *et al.* 2011 | Singapore | 2000–2007 | 34 |
| Hsieh & Chen 2009 | Taiwan | 2007 | 3 |
| Amarakoon *et al.* 2008 | Trinidad and Tobago, Barbados | 1992–2001 | 2 |
| Depradine & Lovell 2004 | Barbados | 1995–2000 | 3 |

*Simulations for excluding coefficients*

As described in the main text, we initially planned to collect data from studies that reported either a correlation between temperature and dengue or a coefficient estimating the effect of temperature on dengue from a regression analysis. However, our systematic literature review revealed that most of the studies using regressions incorporated different covariates into their models, ranging from accounting for no covariates to accounting for the effects of multiple temperature metrics, precipitation, GDP, and others. We therefore conducted a simple simulation to test whether the inclusion of different correlated predictors could result in different estimates for the effect of temperature on dengue.

We simulated a dataset of dengue cases, mean temperature, maximum temperature, and mean precipitation across 10,000 time steps using the rnorm_multi function in the *faux* package in R (DeBruine 2021). In this simulation, the three environmental variables have different means and standard deviations but are strongly correlated with each other and with dengue cases (r = 0.9). We fit five alternative generalized linear models (family=*poisson*) to this same dataset, with predictors including 1) only mean temperature; 2) mean temperature and mean precipitation; 3) mean temperature and mean precipitation and their interaction; 4) mean temperature, mean precipitation, and maximum temperature; 5) mean temperature and maximum temperature. Despite the same underlying data, each of the five models estimated different effects of mean temperature on dengue: model 1 effect = 0.180, model 2 effect = 0.096, model 3 effect = 0.29, model 4 effect = 0.067, model 5 effect = 0.097. The most extreme difference was between model 3 and model 5, in which model 3 estimated an effect of temperature on dengue that is over 4 times larger than model 5.

We repeated the above simulation and model comparison using a weaker correlation between the predictors and dengue and found similar results. Based on these results, we concluded that comparing coefficients from across regression models would be unreliable for the purposes of our study and instead focused on reported correlations from the literature.

**Further details: Does average study temperature impact temperature–dengue effects**?

For this analysis, we aimed to test whether the measured relationship between temperature and dengue depended on the average temperature during the study, as well as whether the correlation between dengue and mean temperature could be accurately predicted from ecological theory based on a lab-parameterized, trait-based model of dengue transmission across temperature (Mordecai *et al.* 2017). We compared a null model and four alternative models, and found that the model that included a quadratic relationship between correlations and average study temperature performed best (lowest AIC). Figure 2 in the main text shows this quadratic model plotted over the residuals of the data, while Figure S1 below shows the same model plotted over the raw data.

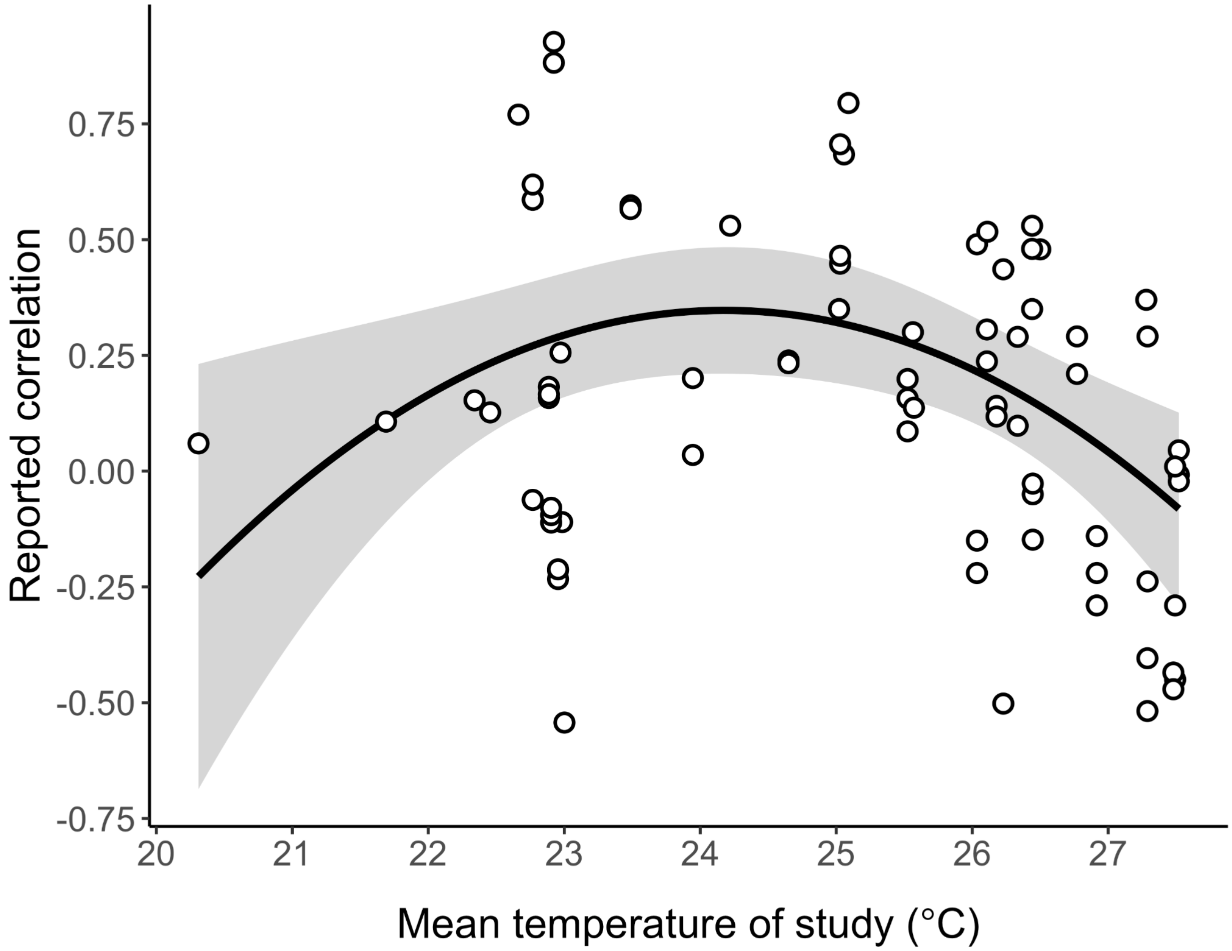


**Figure S1.** Reported correlations between temperature and dengue (points) and fitted predictions (black line) with 95% confidence intervals (shaded region) for the relationship between mean study temperature and reported correlations between temperature and dengue. This model, which included a quadratic effect of mean study temperature (black line), was significantly better than alternative models that included a linear effect or no effect of mean study temperature (ΔAIC from null model = 10.3; pseudo-$R^2$ = 0.209). Confidence intervals were generated using the *effects* package in R (Fox and Weisberg 2019). Figure 2 in the main text shows the same fitted model plotted over the model residuals.

We did not incorporate errors around reported correlation estimates because this information was not available. We repeated the analyses while weighting estimates by the square root of their sample size, a method used in meta-analyses when error estimates are unavailable (Hargreaves *et al.* 2020). The results of this weighted method were qualitatively the same to the unweighted method reported in the main text: the best model included the quadratic relationship between correlations and mean study temperature (ΔAIC between this model and null model = 11.1;

ΔAIC in main analysis = 10.3) and the second-best model included the nonlinear effect of mean study temperature derived from the Mordecai et al. (2017) transmission curve (ΔAIC = 7.7; ΔAIC in main text = 8.0). The model incorporating a linear effect of mean study temperature (ΔAIC = 5.6; ΔAIC in main text = 5.0) did not perform better than the basic model that did not include any effect of mean study temperature (ΔAIC = 6.0; ΔAIC in main text = 5.9).

### Further details: Testing how other climatic and socioeconomic factors explain variation in temperature–dengue effects

The purpose of this section was to test how additional climatic factors such as precipitation, and socioeconomic factors such as country-level GDP impacted the observed effects of temperature on dengue. While we originally intended to estimate how each of these extracted predictors separately mediated the effects of temperature, this was not possible due to the high collinearity between predictors (Fig. S2). We therefore conducted a two step analysis, first collapsing the variance from all predictors with a principal component analysis (PCA), then evaluating the PCA components along with study factors in linear regression models.

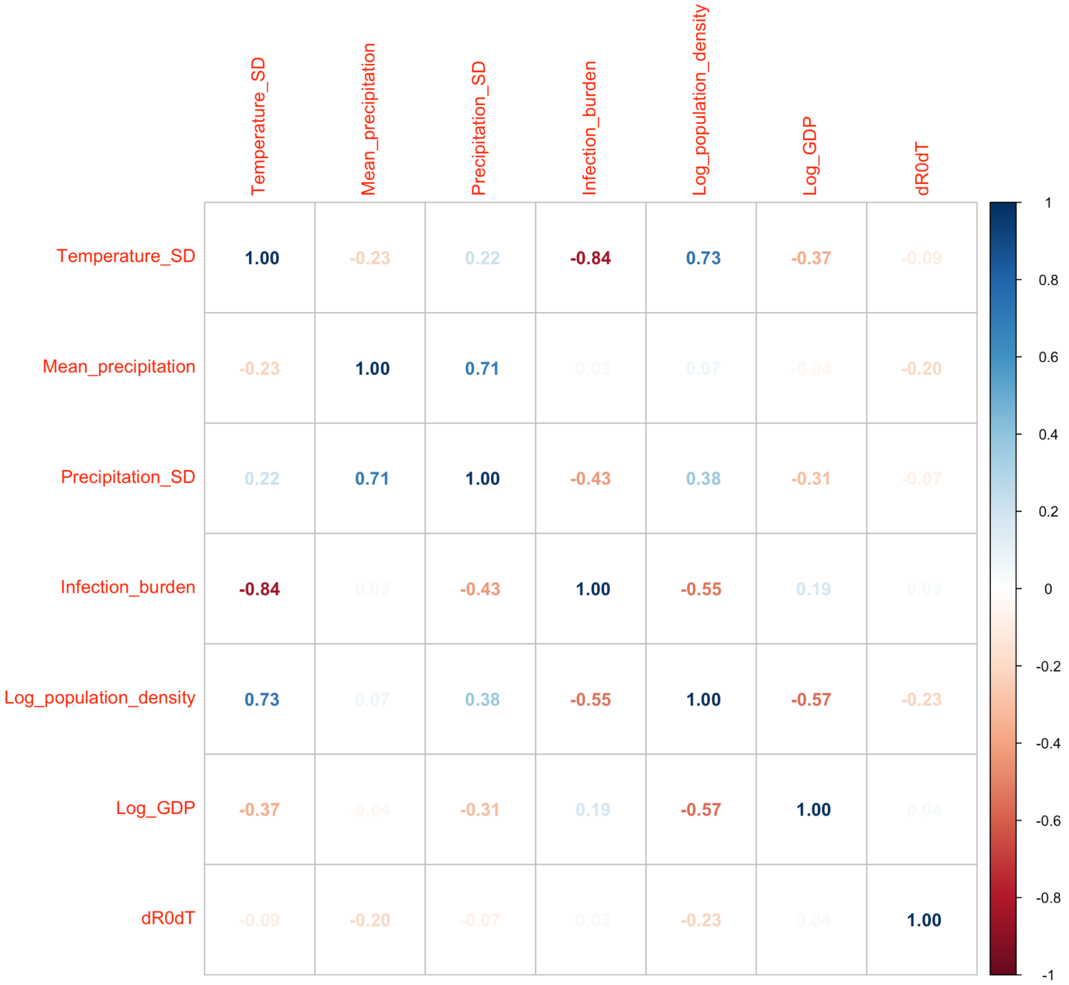


**Figure S2.** Correlation matrix for the seven extracted predictors used in our analysis. Because of high correlations between some of the predictors, we used a two-step PCA and bootstrap regression analysis method.

*Bootstrap regression results: effects of study factors*

The observed effects of temperatures on ecological systems can hinge on the methods and measurements scientists use to detect them. Unlike the climatic and non-climatic mediators primarily discussed in the main text, these study factors affect how effects of temperature are *observed*, rather than the true magnitude of the effects.

The bootstrapped regressions revealed that several study factors had a significant effect on reported relationships. We found that studies using minimum or mean temperature as their temperature metric reported higher correlations compared to those using maximum temperature

(Fig. S3). Additionally, correlations estimated using disease incidence as the response variable were significantly lower than those using cases, and correlations estimated using daily, weekly, or monthly timescales (discrete categories here) were lower than those estimated using annual data (Fig. S3). Correlations did not significantly differ whether they were estimated using Pearson, Spearman's, or autocorrelation methods.

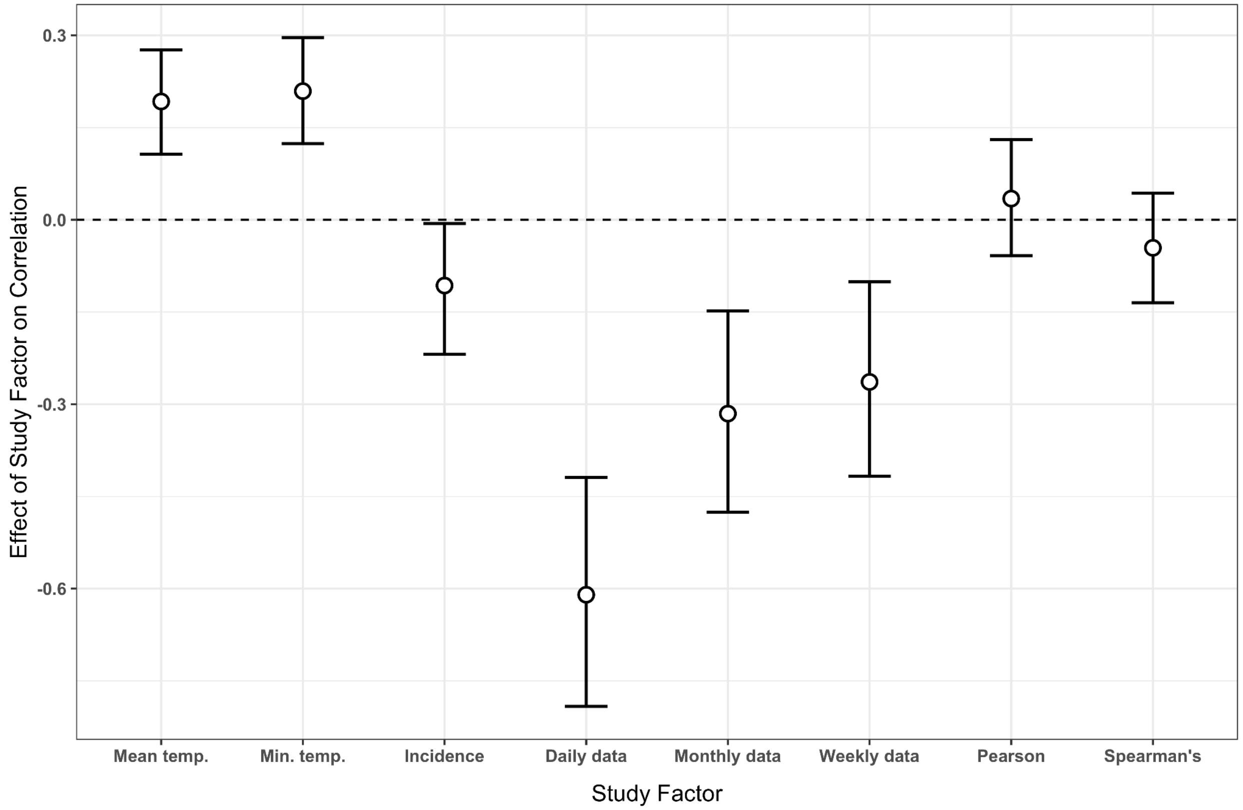


**Figure S3.** Mean and 95% confidence intervals of regression coefficients for the effects of study factors on temperature-dengue correlation across 10,000 bootstrap runs.

The analysis also showed a nonlinear effect of the lag used by a study when modeling a correlation between temperature and dengue, though this tended to be highly variable across the 10,000 bootstraps (Fig. S4). Generally, we found that studies using zero or small lags tended to find slightly negative correlations between temperature and dengue, and that the correlation a study found became positive and stronger with lags between approximately one and four months. The lag results across bootstrap runs are highly variable beyond a four-month lag, likely because our database includes fewer studies using these longer lag periods.

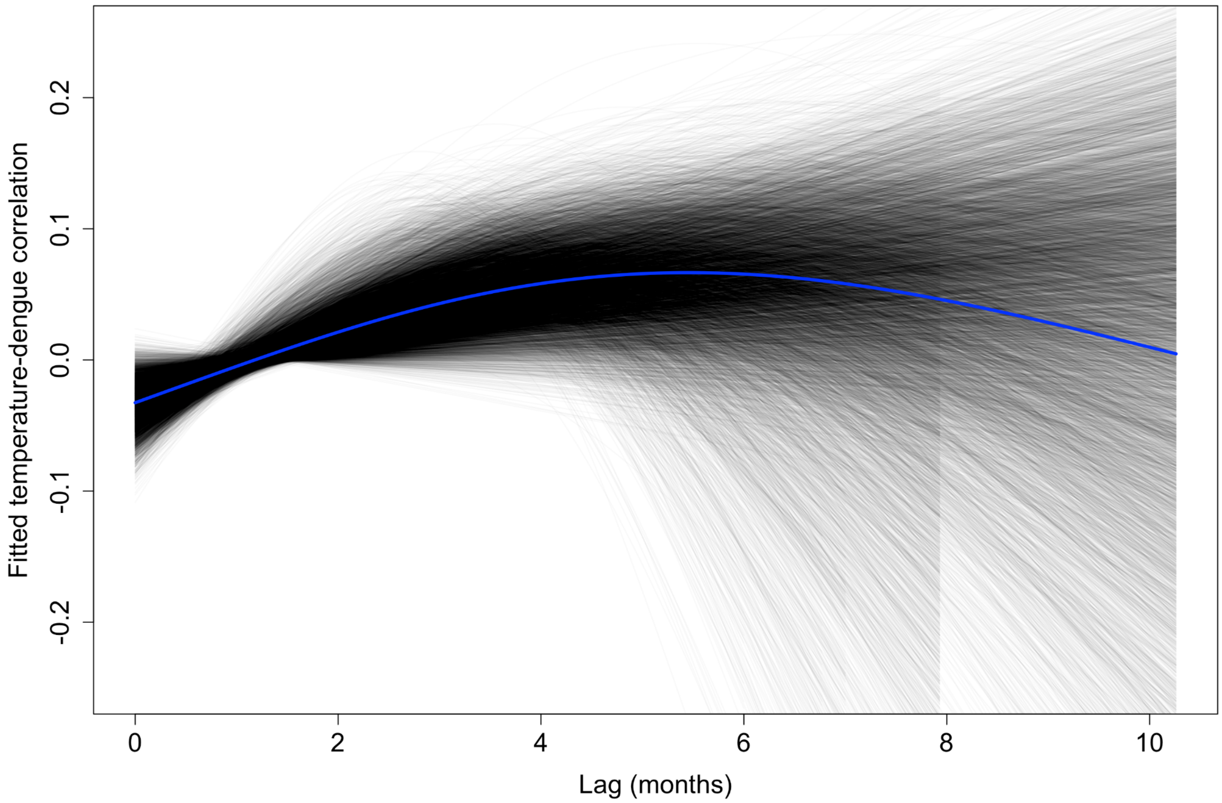


**Figure S4.** Fitted polynomial results for the relationship between temperature–dengue correlations and the lag included by the original studies. Each gray line shows one fit from one of the 10,000 bootstrap runs, and the blue line shows the mean across all runs. Some black lines terminate before others because the longest lag included in a bootstrap run can differ due to the randomly sampled observations included.
